# The divisible load balance problem with shared cost and its application to phylogenetic inference

**DOI:** 10.1101/035840

**Authors:** Constantin Scholl, Kassian Kobert, Tomáš Flouri, Alexandros Stamatakis

## Abstract

Motivated by load balance issues in parallel calculations of the phylogenetic likelihood function, we recently introduced an approximation algorithm for efficiently distributing partitioned alignment data to a given number of CPUs. The goal is to balance the accumulated number of sites per CPU, and, at the same time, to minimize the maximum number of unique partitions per CPU. The approximation algorithm assumes that likelihood calculations on individual alignment sites have identical runtimes and that likelihood calculation times on distinct sites are entirely independent from each other. However, a recently introduced optimization of the phylogenetic likelihood function, the so-called site repeats technique, violates both aforementioned assumptions. To this end, we modify our data distribution algorithm and explore 72 distinct heuristic strategies that take into account the additional restrictions induced by site repeats, to yield a ‘good’ parallel load balance.

Our best heuristic strategy yields a reduction in required arithmetic operations that ranges between 2% and 92% with an average of 62% for all test datasets using 2, 4, 8, 16, 32, and 64 CPUs compared to the original site-repeat-agnostic data distribution algorithm.

## I. INTRODUCTION

Maximizing the efficiency of parallel codes, by distributing the data in such a way as to optimize load balance, represents one of the major challenges in high performance computing.

Here, we address a specific data distribution challenge, which, to the best of our knowledge, has not been addressed before. Our work is motivated by parallel phylogenetic likelihood computations, that is, reconstruction of evolutionary histories based on molecular sequence data (see [1] for an overview). In phylogenetics, we are given a *multiple sequence alignment* (MSA) whose columns are nowadays typically subdivided into distinct partitions (e.g., genes or other disjoint subsets of the data). The columns of a MSA represent the sites and the rows represent the taxa (species) of the MSA for which we intend to infer evolutionary histories. All characters in a given column of the MSA are assumed to share a common evolutionary history. Given the MSA and a partitioning scheme, we can calculate the likelihood on a given candidate tree. In partitioned analyses, individual partitions of a MSA, have a separate set of likelihood model parameters, that is, they are assumed to evolve at different rates. This reflects the distinct evolutionary pressures that, for instance, different genes or parts thereof experience in the course of evolution. Partitioned analyses are the standard use-case for current likelihood-based analyses of empirical data with tools such as RAxaML [2] or MrBayes [3]. In current tools for likelihood-based phylogenetic inference, likelihood calculations on candidate trees typically account for over 85% of total runtime [4]. Thus, the *phylogenetic likelihood function* (PLF) is *the* target function for parallelization. The reasons for load-imbalance in partitioned parallel likelihood calculations are explained in detail in [5].

Initially, we formally state the original data distribution problem. Then, we explain why so-called *site repeats* (SR), that is, sub-columns of the MSA that are identical within a given subtree, complicate the matter.

In the standard case (i.e., without site repeats), we are given a list of *k* partitions and *c* CPUs. Each partition has a computation cost that is linear to the number of sites/columns it comprises. Analogously to the classic bin packing problem, we try to optimally assign the partitions (items) to the CPUs (bins). The first key difference is that the number of CPUs *c* is given, and that we intend to balance the data among all CPUs. In other words, we want to minimize the maximal per-CPU load. The second key difference is that partitions *are* divisible, since a partition consists of (originally independent) sites/subelements. Thus, for improving load balance, we can split partitions into disjoint sets of sites that are allocated to distinct CPUs.

The computational cost per partition has two components: All partitions have an identical constant base cost α. If we split a partition among two or more CPUs, each subset of that partition incurs this base cost α for the CPU it is assigned to. In phylogenomics, α is the cost for calculating the transition probability matrix *P*(*t*) for a given time *t*. This probability matrix is an integral part of the statistical model of evolution. Since partitions evolve under distinct models, we need to compute *P*(*t*) for each partition separately. However, all MSA sites that belong to the same partition have identical model parameters. Thus, α incurs only once per partition and per CPU. Alternatively, we could also parallelize the *P*(*t*) calculations and then broadcast the *P*(*t*) values to all CPUs. However, based on previous computational experiments, the *P*(*t*) calculations are excessively fine-grained and frequent, to allow for efficient parallelization. Therefore, we compute *P*(*t*) (α) redundantly at several CPUs if the sites of a partition have been split up and allocated to more than one CPU.

The second component of the per-partition PLF calculation is the variable cost *φ*. For standard likelihood implementations *φ* is linear in the number of sites per partition. This is because the same amount of arithmetic operations is required to compute individual per-site likelihoods. Since alignment sites are assumed to evolve independently in the likelihood model, calculations on a single site of a partition can be performed independently of and concurrently to all other sites. Therefore, we can easily distribute the sites of a single partition to several CPUs.

Thus, based on the prolegomena, to maximize parallel efficiency, we need to minimize redundant calculations of *P*(*t*) (α) by only splitting partitions when necessary, while distributing sites evenly among CPUs.

Our initial work [6] focused on addressing this data distribution problem for representative PLF implementations in tools such as PLL [7] and RAxML [2]. As mentioned before, the *original data distribution algorithm* (ODDA [6]) assumes that (i) all sites have the same per-site computation cost and (ii) these per-site computation costs do *not* depend on *how* sites of a single partition are split up among CPUs. For example, if a partition with 100 sites is split up among two CPUs and each CPU shall be allocated 50 sites, any split of those 100 sites into 50 sites per CPU will exhibit the same computational cost.

Techniques for accelerating PLF calculations such as *Subtree Equality Vectors* (SEV) [8], [9] or *Site Repeats* (SR) [10], [11] complicate this matter. If we use these techniques in the PLF, the data distribution problem becomes more complex, since both previous assumptions (same cost per site and site independence) regarding per-site computation costs are violated. These techniques take repeating MSA site patterns in subtrees of the phylogeny into account to reduce the amount of per-site PLF computations. A comparison between the PLF functions in the *Phylogenetic Likelihood Library* [7] (derived from RAxML [2]) and a rudimentary implementation that uses SRs shows speedups that range between a factor of 2 up to a factor of 10 [12]. Note that, SR-based techniques can also yield memory savings by more than 50% depending on the input MSA (see e.g., [9]). We describe the SR technique more thoroughly in Section III. Essentially, SR-based optimizations reduce the computation cost *φ* by detecting and re-using identical intermediate PLF results *among* sites. Thus, distinct sites now have varying computation costs. Furthermore, if we assign two sites that can share a large fraction of intermediate results to different CPUs, the accumulated computation cost *φ* will increase. Note that, communicating intermediate shared site computation results between CPUs is not a viable solution, since the computation to communication ratio is unfavorable. Analogously to the argument for recomputating α (*P*(*t*)) we can not communicate such intermediate results because of the extremely fine-grained nature of the PLF operations.

This has implications for distributing data of SR-based PLF implementations. While we can still split partitions arbitrarily among CPUs, we will have to sacrifice some savings that stem from the shared computations. Thus, as for α, we will need to conduct some redundant computations for sites that belong to the same partition and that share some results if these sites are allocated to distinct CPUs. To leverage the substantial computational savings of SR-based PLF implementations, we present an appropriately adapted data distribution algorithm here. Our goal is to split as few partitions among as few CPUs as possible. At the same time we intend to maximize the amount of shared computations between sites for each partition that had to be split. This minimizes the variable per-partition cost *φ* and the accumulated number of base costs α for calculating *P*(*t*). Finally, by reducing the overall accumulated computation cost of the data distribution, we can minimize parallel runtimes.

Given that the problem ODDA aims to solve was shown to be NP-hard, we suspect that the current problem, that has one additional optimality condition, is NP-hard as well. However, a more thorough study of the theoretical properties of our problem is outside the scope of this paper. For this reason, we first assess whether taking into account among-site dependencies for the data distribution algorithm affects parallel efficiency, or not. As we show in Section VI-E, disregarding among-site dependencies *can* decrease parallel efficiency by up to one order of magnitude. In Section V we therefore present and assess several ad hoc heuristics for improving load balance in parallel SR-based PLF computations.

Note that, parallel PLF implementations now form part of several widely-used tools (e.g., ExaML, RAxML, MrBayes, ExaBayes, PhyML) and the results presented here are generally applicable to all of these tools. At present, only ExaML and RAxML offer basic SR-based PLF implementations that merely exploit a fraction of the SRs that are present for accelerating calculations (see [9]). However, we expect that most PLF-based tools will adopt the SR technique [12], because of the substantial runtime *and* memory savings that can be achieved.

The remainder of this paper is organized as follows. In Section II, we survey related work on similar scheduling problems from the area of bin packing with items that can share space. In the subsequent Section III we describe the aspects of the SR technique that are relevant for the work we present here at an abstract level. In Section IV we discuss the properties of the cost factors *α* and *φ*. We also define a theoretical *lower bound* that we use as baseline to design and assess our heuristics. Our main contribution is presented in Sections V and VI. Section V introduces polynomial-time heuristics which yield data distributions that are only 5.75% worse than the lower bound. Finally, in Section VI, we present our experimental setup and the performance gains achieved by our heuristics compared to the original data distribution algorithm (ODDA) on both, simulated, and empirical MSAs.

## II. RELATED WORK

Since we are not aware of any related work in phylogenomics, we briefly review related work on similar scheduling problems.

The so-called *many-constrained bi-objective bin packing* task is one such related problem [13]. Here, items need to be distributed into bins *and* the items are subject to various constraints. The constraints reduce to a cost function between items: The cost of an item can depend on another in a positive, neutral, or negative way if they are in the same bin. In the beginning all items are considered to be located in separate bins. The objective is to minimize the accumulated cost *and* number of bins, by grouping items together into bins.

The Virtual machine (VM) packing problem is also related [14], [15]. The goal is to pack a given amount of VMs (items) into a minimal amount of servers (bins). Each VM has a certain set of memory pages. These memory pages can overlap with those of other VMs, therefore, increasing the number of VMs that fit onto a server.

In contrast to our problem, the items in both scheduling problems are not divisible and the number of bins is not given as input. In fact, the objective function for these related problems is to minimize the number of bins.

## III. SITE REPEATS

Repeated PLF evaluations can account for over 85% of total runtime in maximum likelihood (ML) based tree searches and Bayesian phylogenetic inferences (BI) [4]. One simple observation is that two identical sites that evolve under the same evolutionary model (forming part of the same MSA partition) always yield the same likelihood score. Thus, a common method to avoid these globally redundant calculations is to compress identical MSA sites in a pre-processing step. However, such identical calculations also appear locally at the subtree level. That is, MSA columns might be partially identical for a subset of taxa (species/rows) defined by a subtree (see Figure 1 for an example). While there have been previous attempts to exploit this property [8], Kobert *et al.* recently introduced an efficient algorithm [12] for detecting *site repeats* (SR). The PLF optimization relies on recognizing repeating DNA patterns at different column indices, *i* and *j*, of the MSA (on the same partition), defined by a subtree of the phylogeny whose PLF is being computed. As a consequence, the amount of SR-based savings depends on the actual tree topology.

At an abstract level, PLF calculations conduct post-order tree traversals to update the so-called *conditional likelihood vectors* (CLVs) that have as many entries as the input MSA has columns. Given a fast method to identify repeating subtree site patterns, we can omit all PLF calculations of column indices *p* > *q* for identical subtree site patterns, if *q* is the first occurrence of the specific pattern.

In Figure 1 we only calculate the CLVs at node *v* for column index *p* = 1 and omit the PLF calculations for column indices *q* = 2 and *q* = 5. Without further optimization we can omit a total of 5 out of 15 CLV calculations in this example. An important metric for distributing the data is the number of SRs a specific column contributes to. Henceforth, we will call this metric the *site repeat count* (SRC). In our example site 1 has a SRC of 1, while site 2 has a SRC of 3.

**Figure 1.**
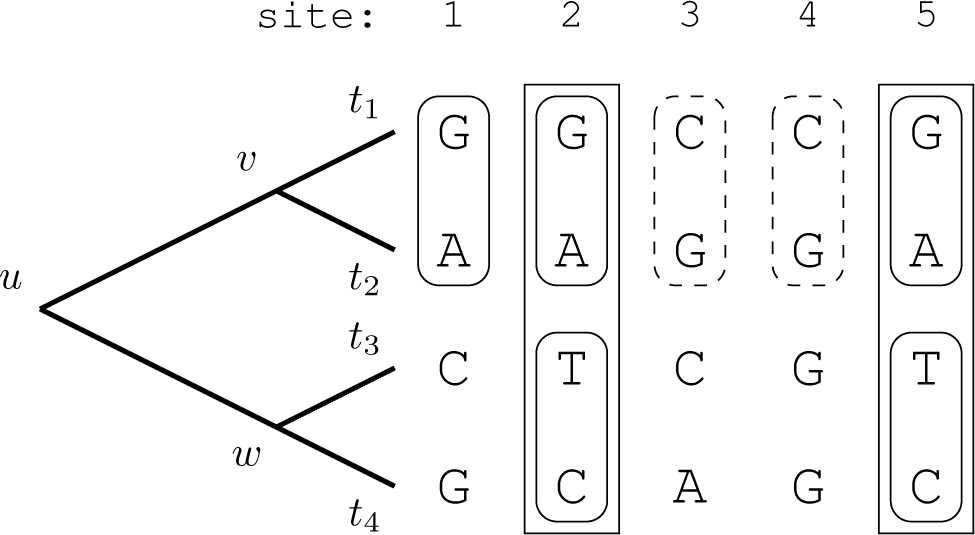
Sites 1, 2, and 5 form a SR at node *v* as they share the same subtree site pattern `GA`. Another repeating pattern is present at sites 3 and 4 (`CG`) for the same node. Note that, node *w* also induces a subtree with pattern `CG` at the tips at site 1. However, since branch lengths can be and typically are different from those of the subtree induced by node *v*, the conditional likelihoods may differ as well. But, there is another SR at sites 2 and 5 for node *w* (`TC`), and hence the conditional likelihood is the same for those two sites. Finally, sites 2 and 5 are a SR for node *u* (`GATC`).

Kobert *et al.* show that a—at a technical level— not fully optimized SR-based PLF implementation consistently outperforms one of the most efficient available PLF implementations (including AVX intrinsics) for distinct real-world application scenarios and tree topologies. Respective speedups range between a factor of 2 up to 10 [12]. Because the work on SRs is very recent, we are not aware of any phylogenomic data distribution algorithm that takes into account the additional constraints induced by SR-based PLF implementations.

## IV. PRELIMINARIES

Assume a MSA of *m* sequences (taxa) and *n* sites 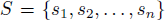, and a partition scheme of *k* disjoint partitions 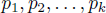 such that 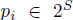. Our task is to distribute these partitions among *c* CPUs, such that the maximum computational cost (*load*) among CPUs is minimized. Note that, we are allowed to assign distinct sites of a partition to different CPUs. Let 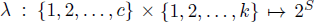 be the mapping of sites to CPUs and partitions. Further, let

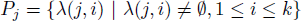

be the set of partitions assigned to CPU *j*. The total computational cost at a CPU *j* is then

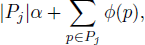

where the mapping 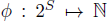 is the SR-based PLF-C (computational cost) for a specific subset of sites from a partition. Note that, for simplicity, in the rest of the text we will treat (site) partitions as lists instead of sets. Hence, we introduce two operations \ and ∪ for adding and removing sites from partition lists. Operation 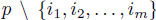 removes sites with indices 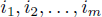 from a partition list *p*, and 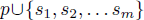 appends sites *s*_1_, *s*_2_,…, *s_m_* at the end of list *p*. A site can be removed in 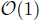 time, and adding *m* sites requires 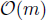 time.

The mapping λ can be implemented as a lookup table of *c* CPUs. Each element of this lookup table is a again a lookup table for each of the *k* partitions. Note that, all algorithms we present assume that the number of sites *n* is larger than the number of CPUs *c*.

### A. Computational costs α and φ

The first simple observation is, that the computational cost for each CPU depends on the cost components α and *φ*. However, both quantities can only be measured in abstract terms. The exact runtimes depend on the hardware architecture, data type, and tree topology. Therefore, the objective is to minimize both factors, without being able to directly compare them. We will show that the count of additional α values, which we call *extra*_α_, created by splitting partitions, does not vary substantially for the heuristics we tested in Section VI-E. Thus, we focus on minimizing the cost *φ*, which corresponds to the number of arithmetic operations required to evaluate the PLF on a given set of sites for a given tree topology. We call this cost the *PLF count of operations* (PLF-C).

### B. Lower bound *L*

The second observation is, that we can calculate a simple theoretical lower bound for the optimal accumulated per-CPU PLF-C. We use this bound, to steer our heuristics *and* to assess the performance of our heuristics.

Assume that all partitions are assigned to a single CPU. We can then calculate the PLF-C for this CPU. Simply dividing this single CPU PLF-C by the number of CPUs *c* yields a natural lower bound *L*. Since no partitions have been split up on the single CPU assignment, the respective SR-induced savings are maximal. Thus, for any assignment of the partitions to two or more CPUs, which may not be able to retain all site repeats, the per-CPU PLF-C will be ≥ *L*. Consider the example MSA in Figure 1 with only two sites and assume that we want to assign the data to two CPUs. The lower bound is 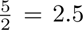 PLF-C. However, the optimum data distribution will assign one site to each CPU. Thus, the optimized PLF can not skip the SR `GA` anymore, resulting in a per-CPU PLF-C of 3 > 2.5 for both CPUs.

## V. DESCRIPTION OF HEURISTIC COMPONENTS

The heuristics we present here consist of three phases: Data pre-processing (phase I), initial distribution of sites to CPUs (phase II), and site reshuffling strategies (phase III). The data pre-processing phase prepares the MSA data for the subsequent initial distribution phase. We present two algorithms for computing such an initial partition data to CPU distribution. Finally, we employ three different site reshuffling strategies to further decrease the PLF-C.

The individual components designed for these three phases can be flexibly combined in a plethora of ways and orders to rapidly assemble and test heuristic data distribution algorithms. We assessed a total of 72 heuristic strategies using the 7 core components (2 for pre-processing, 2 for the initial assignment, and 3 for the reshuffling phase) for implementing the three distinct distribution phases. For a complete list of all 72 strategies, see the on-line supplement^1^ on `github`. Here, we only present a performance assessment of 32—including the two best-performing ones— out of 72 heuristics. Presenting all 72 heuristics would merely require describing additional variants of our components.

In addition, we describe the two heuristics that performed best, in more detail. All 32 heuristics we tested are enumerated on the *x*-axis of Figure 12.

**Figure 12.**
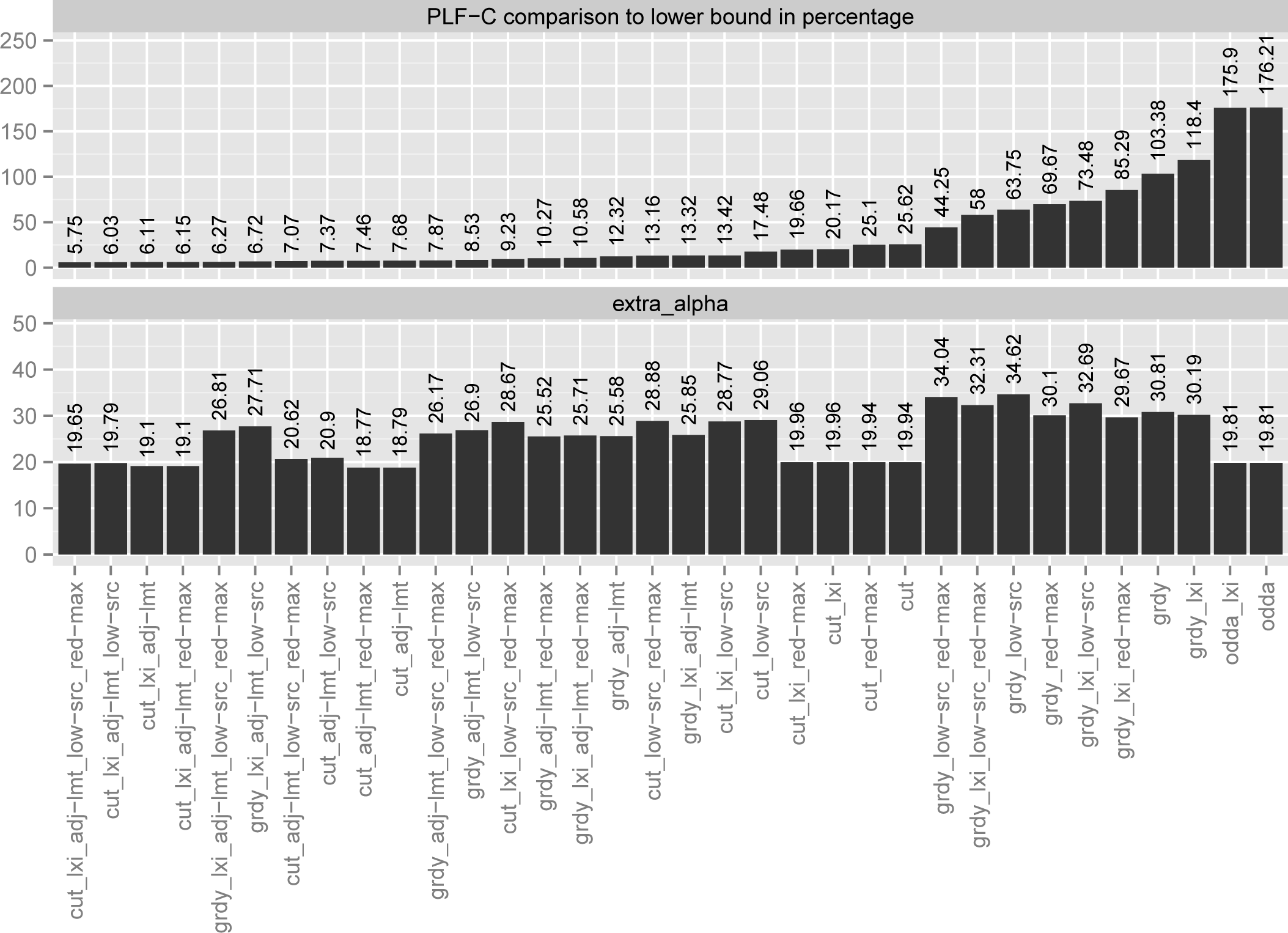
Comparison of all 32 heuristic component combinations, where, cut = cut approach; grdy = greedy approach; lxi = lexicographic sorting of MSA; adj-lmt = adjust limit reshuffling strategy; low-src = low SRC reshuffling strategy; red-max = reduce max reshuffling strategy. The upper part presents the average PLF-C to *L* percentage of the heuristics over all MSAs and numbers of CPUs. The lower part presents the average *extra_α_* for the heuristics over all MSAs and numbers of CPUs.

### A. Phase I heuristics: Data pre-processing

A commonly used method to avoid redundant calculations in the PLF is to remove all duplicate sites from the MSA and only keep the unique sites. Since identical sites yield the same per-site likelihood if they evolve under the same model, the overall likelihood can be computed by assigning respective weights to unique sites.

As a second pre-processing step we sort the remaining, unique sites lexicographically. Through an empirical analysis we have observed that lexicographic sorting has the effect that sites sharing repeats are more likely to reside next to each other than for a random site order. Figure 10 depicts the sorted versus unsorted PLF-C between all pairs of consecutive sites, accumulated over all sites, all 1200 inferred trees, and all 8 MSAs we used (see Section VI-B). Sorting leads to better results for the overall data distribution algorithm (see Section VI-E). The lexicographic sort requires 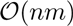 time and 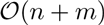 space.

**Figure 10.**
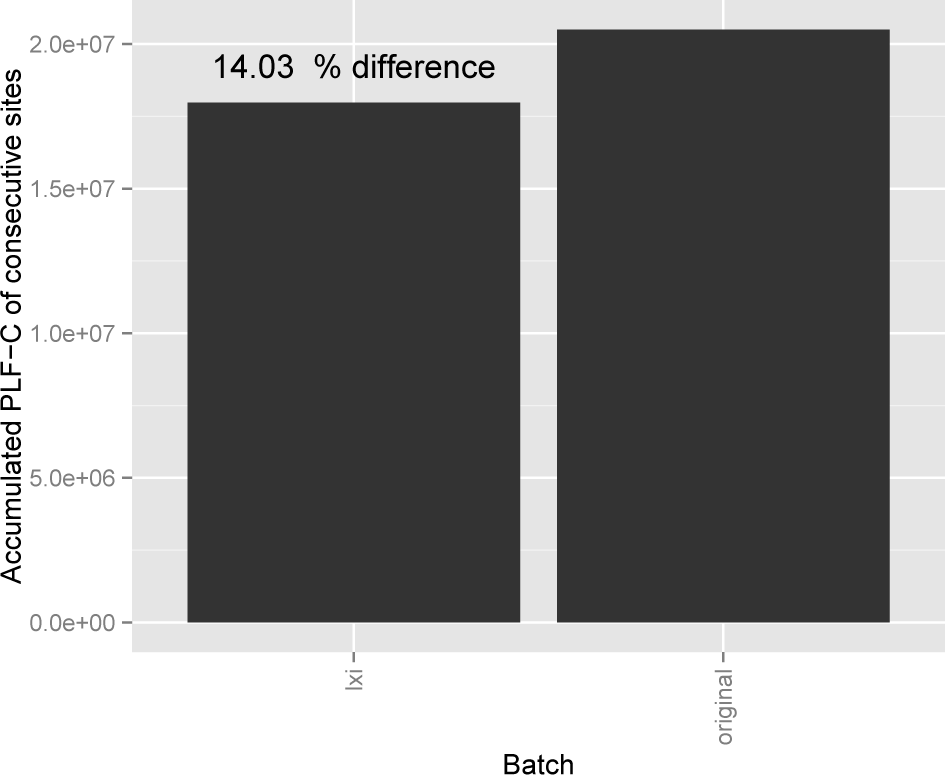
Lexicographically sorted versus unsorted PLF-C between all pairs of consecutive sites, accumulated over all sites, all 1200 trees, and all 8 MSAs.

### B. Phase II heuristics: Initial distribution

The initial distribution of partitions to CPUs is a two-step process. First, we *pre-fill* the CPUs with entire partitions, that is, without splitting any partition until a filling limit is reached for at least one CPU. Then, we distribute and potentially split up the remaining partitions via the *greedy* or the *cut* approach (see below). Both approaches try to balance the PLF-C among CPUs.

In the pre-filling step, we intend to assign as many entire (without splitting them up) partitions as possible to the CPUs, for minimizing the per-CPU α cost. Initially, the *lower bound L* (see Section IV-B) is used as a computing capacity limit for each CPU. The reshuffling strategies (phase III, see Section V-C) might adjust this limit later-on. The procedure for assigning entire partitions to CPUs is analogous to phases 1 and 2 of the ODDA [6]: We first sort partitions by increasing order of computational cost (PLF-C). Starting with the partition that has the smallest PLF-C, we assign partitions to CPUs in a cyclic manner until adding a partition exceeds the capacity limit (*L*) of a CPU. When the procedure stops, the still unassigned (remaining) partitions have indices 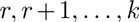. In a final step we sort the CPUs by their accumulated PLF-C in ascending order. Figures 9a and 9b illustrate the pre-filling process; Figure 2 contains the PREFILL algorithm. The asymptotic runtime complexity for pre-filling is 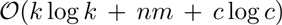, as sorting the *k* partitions requires time 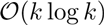, and calculating the PLF-C for all partitions requires 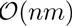 time.

**Figure 2.**
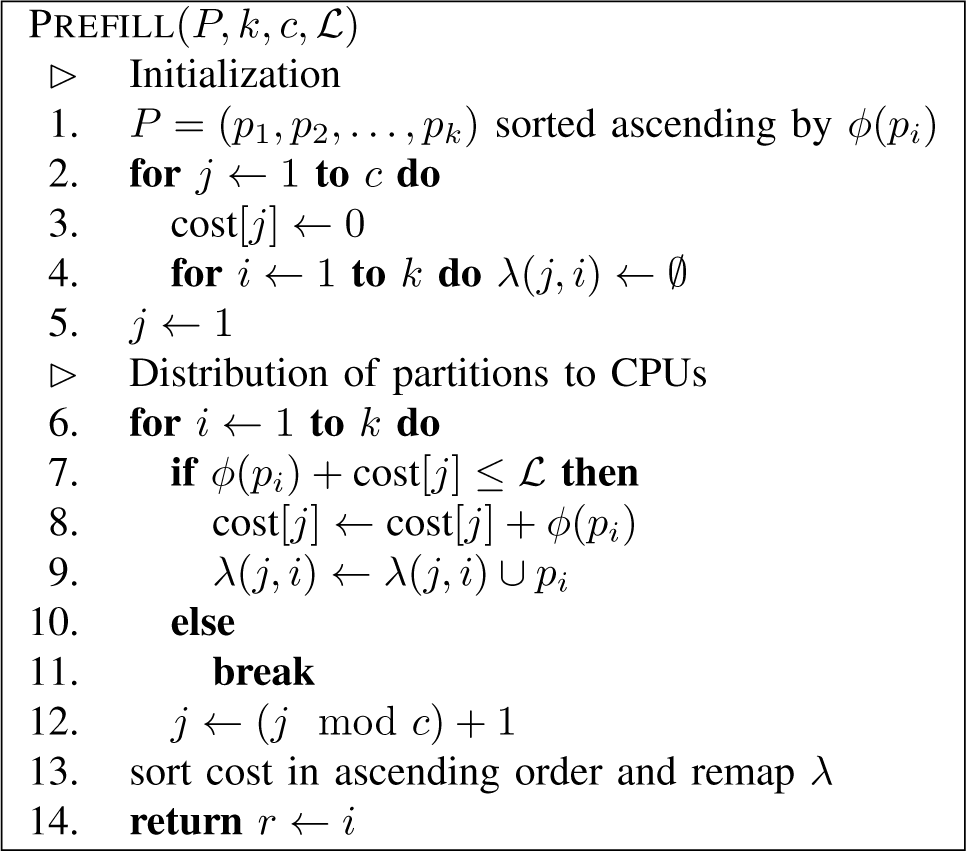
The PREFILL function requires the following input parameters: a list *P* of *k* partitions sorted in ascending order by their PLF-C, the number of CPUs *c*, and the lower bound *L*.

1) *Greedy approach:* The greedy approach is one of two alternatives for distributing the remaining partitions among the pre-filled CPUs. It tries to find the best fit CPU for each individual site. The site fit to a CPU is defined as the PLF-C increase that is induced by assigning the site to the CPU. The lower this PLF-C increase is for a specific CPU, the better the site fits. The underlying idea is, that by increasing the fit of sites from remaining partitions that need to be split up we can reduce the overall PLF-C for the data distribution.

However, we need to first assign at least some sites of the remaining partitions to CPUs to be able to compute a meaningful site fit. We achieve this via a virtual assignment of remaining partitions to CPUs (please see Figures 9c-9f before reading on, since the subsequent description will be easier to follow). Initially, we calculate the average per-site PLF-C cost μ for each remaining partition. Then, we use μ as a proxy for the PLF-C to generate a virtual assignment of the remaining partitions to the CPUs. This virtual assignment ignores SR effects and simply distributes all sites from all remaining partitions to the CPUs based on μ. Thereby, we can approximately balance the per-CPU PLF-C. Then, for each CPU that has at least one site from one of the remaining partitions, we select the site in the middle of the subset of sites of the remaining partition (the median site) that has been assigned to this CPU as representative site. This representative site from the virtual assignment is then assigned to the respective CPU and becomes a real site assignment. As a result, at most two sites from at most two of the remaining partitions are assigned to each CPU. Sites of at most two distinct remaining partitions are assigned to each CPU, since we virtually assign the remaining partitions one-by-one in as large as possible monolithic blocks to CPUs. Keep in mind that each of the remaining partitions exceeds the capacity limit of every CPU because of the way we designed the prefilling component.

Figures 9c-9e illustrate and Figure 3 presents the pseudocode for the virtual assignment step of GREEDY. In the pseudocode *z* is the site index pointing to the first site of the current partition, while *z*’ is the length of the current partition. The variable s in line 9 is the median site. Since we remove the median site s from *p_i_* we also need to decrement *z* for the next partition (line 13). If the algorithm virtually assigns two partitions to a CPU, *vcost* represents the computational cost that is required for the first partition that was assigned to this CPU and defines how much computing capacity is left for the second partition on this CPU.

**Figure 3.**
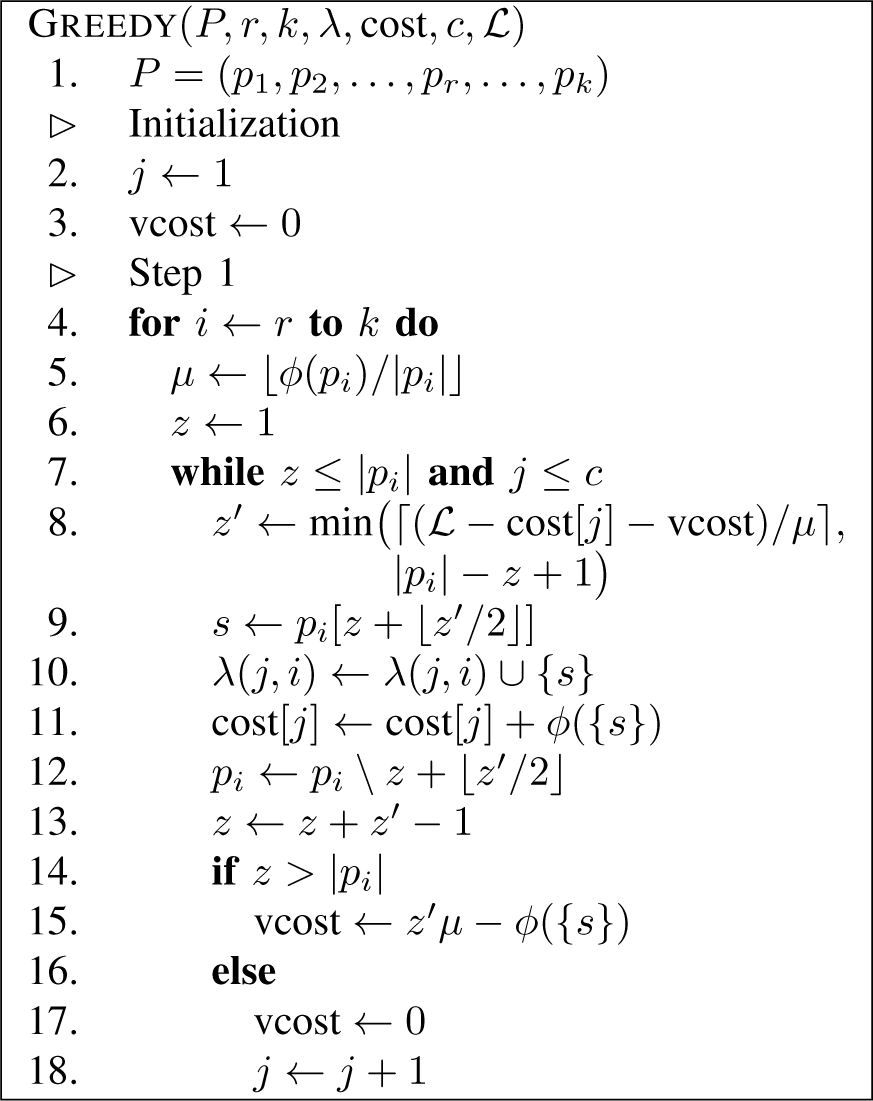
The GREEDY function requires the following input parameters: The list *P* of remaining partitions sorted in ascending order by their PLF-C, where index *r* denotes the first and index *k* the last unassigned partition; the distribution function λ and the *cost* for the *c* CPUs; the lower bound *L*. Here we show the virtual assignment step (step 1) of the algorithm.

Once we have computed this initial assignment of median sites from the remaining partitions to CPUs, the second step of GREEDY is straight-forward. We simply iterate over the yet unassigned sites from the remaining partitions and assign each site to the CPU with the best PLF-C fit. If adding a remaining site to this CPU leads to exceeding the capacity limit *L*, we assign the site to the CPU with the second-best PLF-C fit etc., provided that adding the site will not violate its capacity limit. As a consequence we generally assign the site to a CPU that already has some sites of the same partition in order to minimize the α cost. Only, if no CPU below the capacity limit has sites of the same partition, but there are other CPUs below the capacity limit, we will assign the site arbitrarily to one of the latter. Finally, if *all* CPUs exceed *L*, we assign the site to the CPU with the best fit. Figure 4 presents the pseudocode for the second step of GREEDY. Note that, the second step of Greedy will only terminate once all sites of all remaining partitions have been assigned to a CPU.

**Figure 4.**
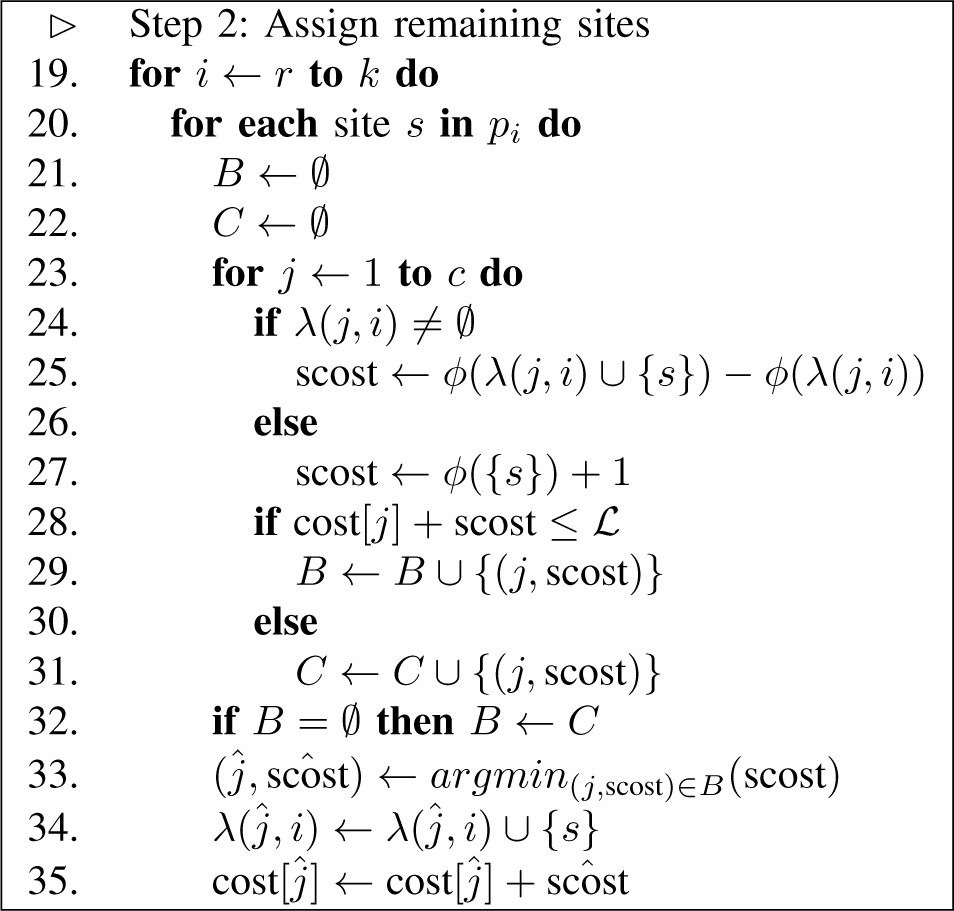
Step 2 of the GREEDY function.

The total asymptotic runtime of step 1 is 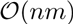. The runtime is dominated by calculating μ in line 5. For all sites in the *r – k* remaining partitions we have to compute φ, which requires 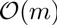 time per site.

The runtime of step 2 is 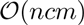. For each site of the *k – r* remaining partitions we need to calculate the *scost* for each CPU. The *scost* in line 25 can be computed in 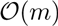 by keeping the site repeats data structure in memory and using the algorithms from [11], [16]. We use this observation in all subsequent runtime analyses.

*2) Cut approach:* As an alternative to GREEDY, we implemented the CUT approach for distributing the remaining partitions to pre-filled CPUs. Here, we cut contiguous regions from the remaining partitions and assign them to CPUs. This has the benefit of incurring *exactly* the same accumulated α cost as the ODDA, which has been proven to exceed the optimal solution by at most one α per CPU [6]. We illustrate the cut approach in Figures 9g and 9h.

After the pre-filling phase, the remaining partitions and CPUs are still sorted by increasing PLF-C. Starting with these sorted lists, we iterate over the CPUs from the least filled to the most filled one (i.e., from largest to smallest available capacity). For each CPU we select a region from at most two of the remaining partitions. The size of the region is computed as follows for every CPU to be filled: we divide the free capacity of the particular CPU by the accumulated free capacity of all CPUs that have not reached their capacity limit yet and multiply this by the total PLF-C of all remaining and yet unassigned partitions. The underlying idea is to distribute the additional arithmetic operations we will need to conduct due to partitions that have already been split up as evenly as possible over all CPUs. To then actually assign such a cut region to the CPU, the region must exhaust or exceed the capacity limit of the current CPU. If this is not the case, we increase the size of the region until it exhausts the capacity limit of the current CPU. This restriction that CPU capacity limits must at least be exhausted is required for the reshuffling strategies (phase III, see V-C).

The pseudocode for CUT is provided in Figure 5. While δ returns the free capacity for a given CPU *j*, the array Δ contains the accumulated free capacity of all CPUs with an index ≥ *j* for a given *j*. We use *z* as site index again and *rcost* is the total PLF-C of all remaining partitions. The value *l* denotes the target PLF-C capacity of the current CPU *j* which defines the size of the region that shall be assigned to CPU *j*. Note that, CUT terminates when all sites from all remaining partitions have been assigned to CPUs.

**Figure 5.**
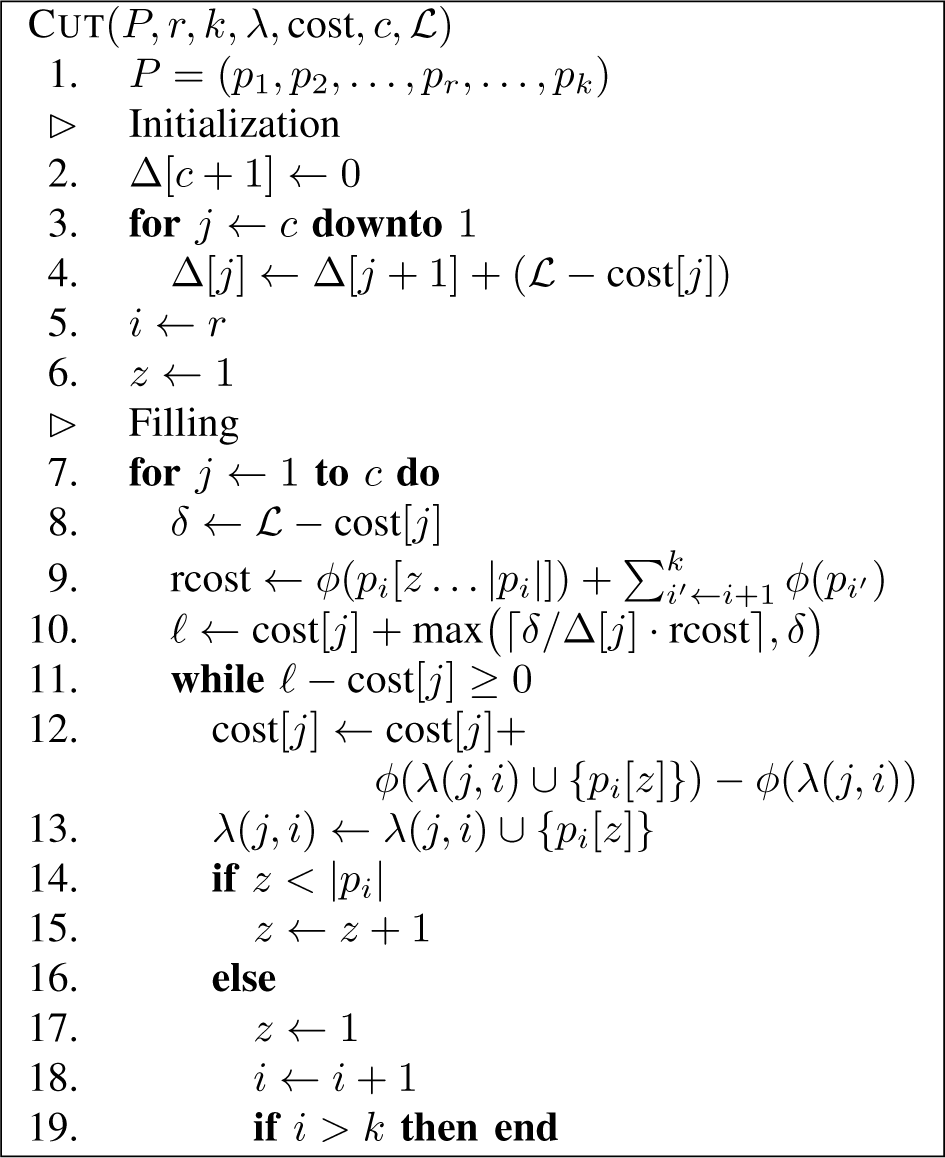
The CUT function requires the following input parameters: The list *P* of remaining partitions sorted in ascending order by their PLF-C, where index *r* denotes the first and index *k* the last unassigned partition; the distribution function λ and the *cost* for the *c* number of CPUs; the lower bound *L*.

The total asymptotic runtime of CUT is 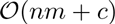. For line 9 we can initialize two arrays: One, computing *φ* in the order of sites in the remaining partitions and the other computing *φ* in the reverse order of sites. Thus, line 9 requires constant time and the initialization can be computed in 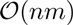. By definition we will split up at most 2(*k – r*) partitions. These partitions are disjoint. Therefore, line 12 will be executed once for each site of the partitions that will be split and thus requires 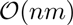 time.

### C. Phase III heuristics: Reshuffling strategies

After the initial distribution (e.g., computed with GREEDY or CUT), we can employ up to three site reshuffling strategies to further improve the PLF-C: Adjust limit, reduce max, and low SRC.

1) *Adjust limit:* The initial distribution provides a good estimate of how many partitions need to be split up and how the overall PLF-C for the distribution increases due to these splits. We use this knowledge to adjust the capacity limit of the CPUs: Instead of the lower bound *L* we now use the average CPU load from the initial distribution as capacity limit. We then simply re-run the pre-filling and the GREEDY or CUT approach with this updated capacity limit.

2) *Reduce max:* The bottleneck in parallel PLF computations is always the CPU with the highest PLF-C since it determines the overall runtime. Therefore, we attempt to reduce the load of the CPU with the highest PLF-C.

This reshuffling strategy uses the worst-case PLF-C for one site. This is a MSA-specific value that depends on the number of taxa *m*. The *worst-case site PLF-C* simply has a SRC of zero.

First, we select all partitions that have been split up and have been assigned to the most loaded CPU. Then, we determine all candidate CPUs that also have a part of those partitions. Out of those candidate CPUs we then select only those with a PLF-C that is smaller than the mean per-CPU PLF-C. Using the worst-case site PLF-C, we estimate how many sites can be moved from the most loaded CPU to the least loaded CPUs, such that they do not exceed the current capacity limit. We then move the sites with the lowest SRC away from the most loaded CPU. Thereby, we only lose a small number of SRs for the sites that will remain on the most loaded CPU. This *reduce max* operation can be repeated for an arbitrary number of times. We choose to repeat this process as many times as there are CPUs to achieve an acceptable trade-off between overall PLF-C improvement and the run-time for this heuristic component. Evidently, one could also design an explicit convergence criterion to determine when additional applications of reduce max are not worthwhile any more.

We present the pseudocode for reduce max in Figure 6. Line 1 assumes that *P* is the original sorted list of all partitions from the pre-filling phase (Section V-B). The constant *wcost* is the *worst case PLF-C for a site. P’* is the list of split up partitions assigned to one of the most loaded CPU *j*.

**Figure 6.**
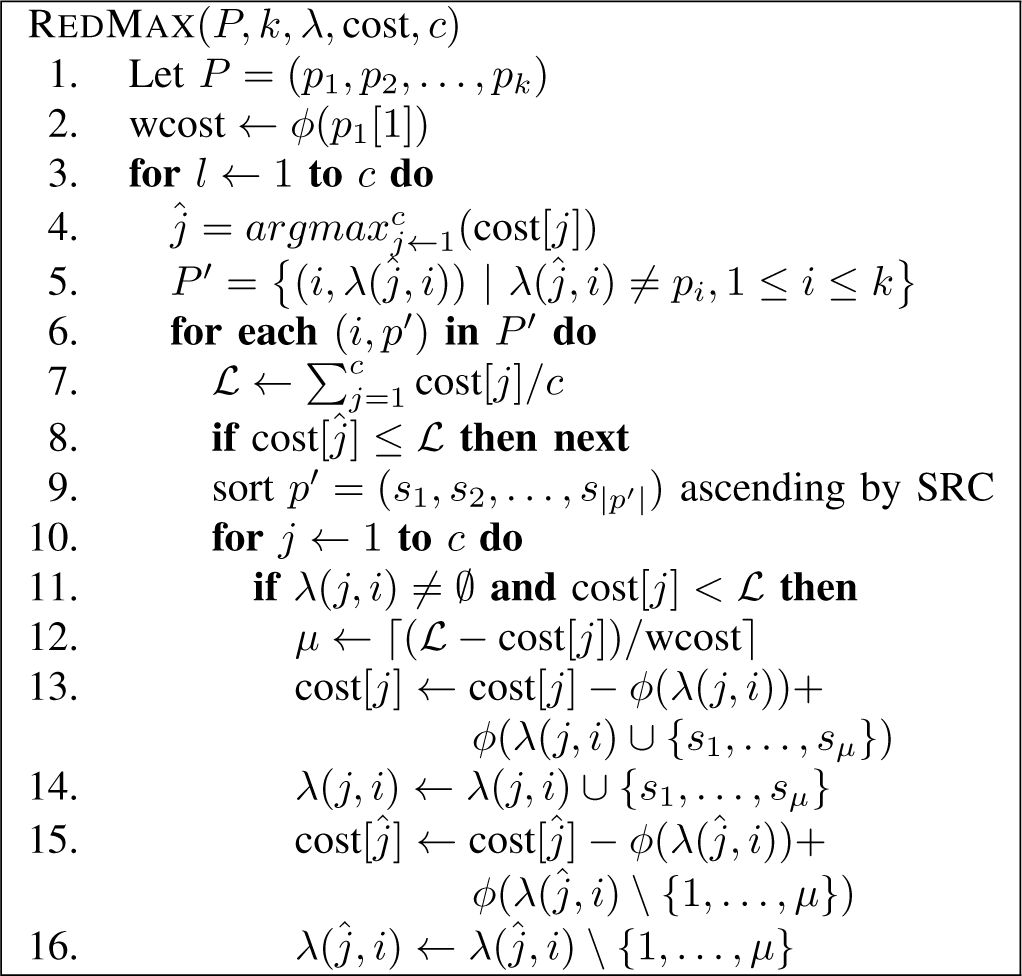
The REDMAX function requires the following input parameters: a list *P* of *k* partitions, the distribution function λ, and the *cost* for *c* CPUs.

The total asymptotic runtime for *reduce max* is 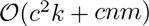. The first part is defined by the calculation of *L*. The second part is defined by sorting the sites in *p*’ by their SRC.

3) *Low SRC:* One goal of our distribution strategies is to maximize the site repeats in order to decrease the overall PLF-C. The *low SRC* strategy intends to reshuffle sites that exhibit a low SRC on their current CPU. For reshuffling, we select 20% (see below for a rationale for this setting) of the sites from each split up partition that have the lowest SRC under the current data distribution. We then provide this selection of sites as input to step 2 of GREEDY (see Section V-B1) to reassign them to the CPUs.

The actual percentage of sites selected for reshuffling is a tuning parameter. It should not be too high, since we want to determine how well the sites fit into a comparatively large, fixed site distribution. It should not be too low either, since this will decrease the overall PLF-C improvement of the reshuffling. We empirically determined that a threshold setting of 20% yields good performance.

Figure 7 presents the pseudocode. Line 1 assumes that *P* is the original sorted list of all partitions as obtained in the pre-filling step (Section V-B). *P*’ is the list of split partitions of CPU *j*.

The total asymptotic runtime of *low SRC* is 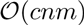. For each CPU we iterate over all split partitions, which are at most *k* and sort them in 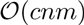. Line 9 will reassign at most *n* sites, where each site requires 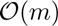. As before Greedy phase 2 requires 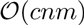 time.

**Figure 7.**
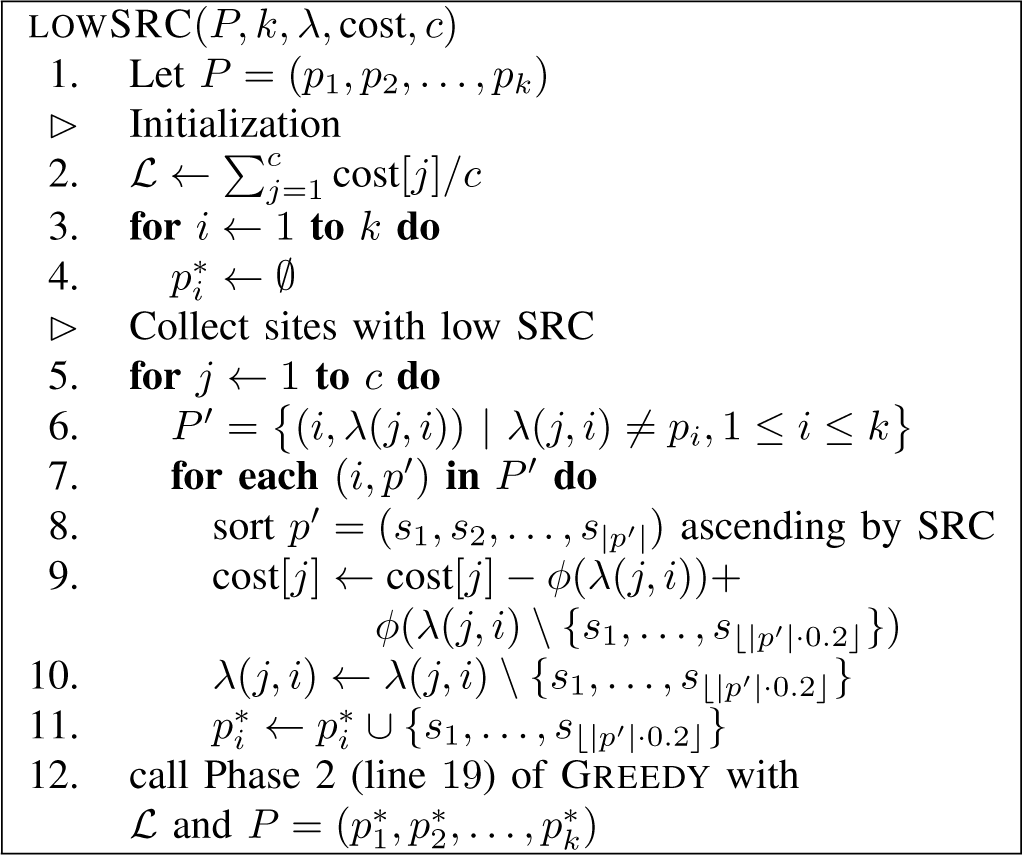
LOWSRC takes the following input parameters: a list *P* of *k* partitions, the distribution function λ, the *cost*, and the number of CPUs *c*.

## VI. EXPERIMENTAL SETUP AND RESULTS

### A. Software

Given a MSA and a partition scheme, we employed RAxML [2] to infer and sample phylogenetic trees (for details see Section VI-C). Using Ruby and the Ruby Gem Newick-Ruby^2^ for parsing trees, we initially characterized the problem and then implemented the heuristics. To visualize and post-process the results we used appropriate `R` packages.

### B. Test datasets

We used five simulated and three empirical nucleotide MSAs for benchmarking the heuristics. To simplify the experiments we removed all identical sites from the MSAs prior to running the heuristics such that all MSAs only contained unique sites. The MSA are available for download at https://github.cormconscho/phylo_scheduling/ttee/master/ datasets and their properties are summarized in Table I.

**Table I.** MSA PROPERTIES.

| ID | Type | Taxa | Unique Sites | Partitions |
| --- | --- | --- | --- | --- |
| 1 | Empirical | 59 | 3238 | 8 |
| 2 | Empirical | 128 | 19,866 | 34 |
| 3 | Empirical | 404 | 7444 | 11 |
| 4 | Simulated | 10 | 76,754 | 100 |
| 5 | Simulated | 10 | 169,939 | 200 |
| 6 | Simulated | 10 | 199,974 | 232 |
| 7 | Simulated | 10 | 246,897 | 310 |
| 8 | Simulated | 10 | 269,697 | 332 |

### C. Characterizing the problem

Initially, we assessed to which degree the actual tree shape influences the PLF-C differences between SR-based and SR-agnostic PLF implementations. We started 100 maximum likelihood (ML) tree searches from randomized stepwise addition order parsimony starting trees and another 100 ML searches on random starting trees for every test MSA using RAxML. We then calculated the PLF-C ratios between SR-based and SR-agnostic PLF implementations for each tree and MSA. The average SR-induced PLF-C savings for ML trees inferred on random and parsimony starting trees are almost identical. However, ML trees inferred from random starting trees show a slightly higher variance in savings than ML trees inferred on parsimony starting trees. This variance is more pronounced for our empirical MSAs. Figure 11 shows a representative boxplot for MSA 3.

**Figure 11.**
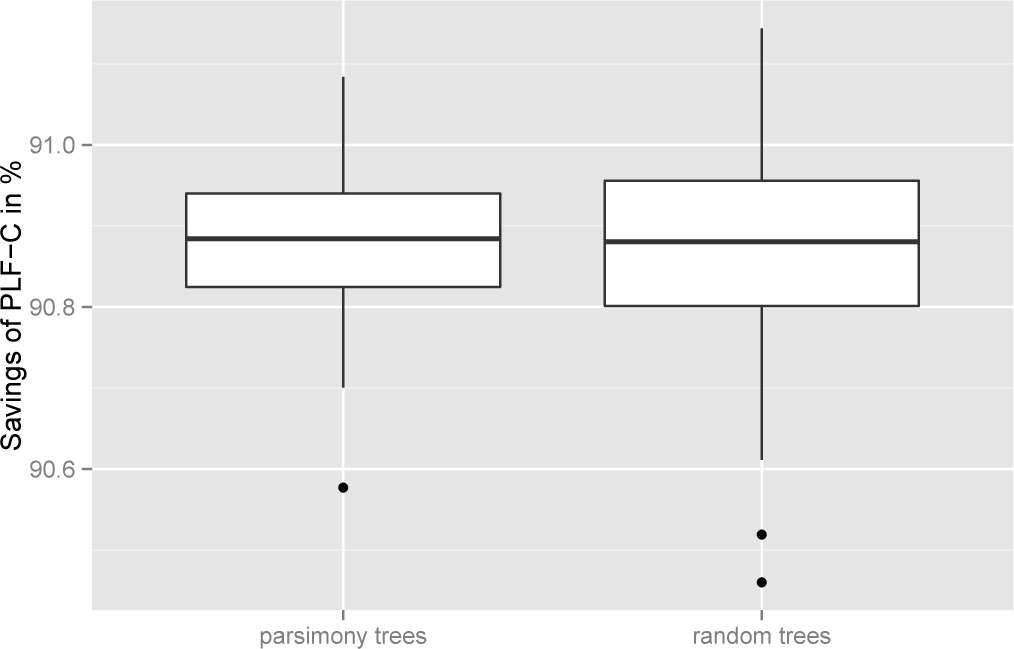
PLF-C savings when employing the SR technique for MSA 3

As shown by Kobert *et al.* [12], the savings are substantial. The average PLF-C saving over all MSA and all trees we generated is 61.68%, with a minimum of 48.92% and a maximum of 91.14%.

Next, we evaluated possible factors influencing the SR-induced savings based on several MSA characteristics. We did not find a correlation of savings with the likelihood score of a tree for a given MSA. That is, a tree with a better likelihood score does not necessarily have a lower PLF-C. Also, the PLF-C does not appear to correlate with the number of taxa. However, the number of sites per partition *is* positively correlated with PLF-C savings. Also, the placement of the virtual root, that is used to conduct the post-order tree traversal for calculating the CLVs (see Section III), has an impact on the PLF-C. Our initial experiments revealed that placing the virtual root using the so-called midpoint rooting technique, that reduces the height of the thereby *rooted* binary tree, increases PLF-C savings.

Because of this observation, all of our heuristics use a midpoint rooting on the given input tree for calculating the PLF-C of respective data distributions.

### D. Ground-truth comparison

To analyze the performance of our heuristics we also compared them with the exact, optimal solution. We therefore generated and evaluated all possible data distributions exhaustively. Since there are n^c^ possibilities for assigning sites to CPUs, this was only feasible for small problem instances. The optimal solution is the data distribution that minimizes the PLF-C of the most loaded CPU. From each of the 8 MSAs in Table I we assembled 4 small MSAs by sampling MSA sites from the beginning of each partition. These groundtruth MSAs contained 4 partitions with 3 – 5 sites that were assigned to 2 CPUs and 2 partitions with 4-6 sites that were assigned to 3 CPUs. As one may expect, the exact solution is generally better than our heuristic solution. However, for 11 out of the 32 MSAs one of the heuristics was *on par* with the exact solution. In the worst case, the PLF-C difference between the best heuristic and the optimal solution for the most loaded CPU was 9.34%.

The optimal solution was also the *perfect* solution in 1 out of 32 experiments. We have a perfect solution when the PLF-C of each CPU equals the lower bound *L* (see Section IV-B). For instance, this is the case if the data distribution does not need to split up any partition such that sites which share SRs are not distributed over different CPUs (see Section III).

Keep in mind that, due to the small input sizes, the above results are mainly useful for verifying that our heuristics are not ‘far off’. Also note that, the groundtruth calculates the optimal solution with respect to *φ* (i.e., the PLF-C) but does not strive to minimize the α cost.

### E. Performance of heuristics

To assess the performance of our heuristics, we calculated data distributions for all MSAs. For each MSA we arbitrarily selected one of the ML trees inferred on a parsimony starting tree with RAxML and computed distributions for 2, 4, 8, 16, 32, and 64 CPUs. We then averaged the results for each heuristic over all MSAs and CPU counts. We assess the resulting heuristic data distributions by comparing the respective PLF-C of the most loaded CPU with the lower bound PLF-C L (see Section IV-B) and the resulting number *extra_α_* of additional *P*(*t*) calculations due to splitting up partitions (where 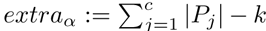, see Section IV for notation *c*, *P_j_*, and *k*) with the lowest *extra_α_* among all heuristic distributions. The heuristic that yielded the lowest PLF-C over all MSAs comprises the following components: a lexicographic MSA sort, then CUT, followed by the ADJUSTLIMIT, LOWSRC, and REDMAX reshuffling strategies. On average, the PLF-C of this heuristic is only 5.75% higher than the rather conservative theoretical lower bound *L*. Out of the 32 combinations of our heuristic components that we list in this paper (see Section V) it ranks 5*th* with respect to *extra_α_*. The *extra_α_* is only 4.69% higher than that of the best result which we deem acceptable based on our experience with developing ExaML and ExaBayes.

The best combination using the GREEDY component, also uses lexicographically sorted MSAs and *all* available reshuffling strategies in the same order as above. Its PLF-C is on average 6.27% worse than *L* but the *extra_α_* is 42.83% higher than that of the best result which might be rather prohibitive for real-world parallel PLF implementations.

We also need to answer the question if a dedicated data distribution strategy for SR-based parallel PLF implementations is indeed necessary. To this end, we repeated the above experiments with the original SR-agnostic data distribution algorithm (ODDA). The data distribution proposed by ODDA requires on average 176.21% more PLF-C than *L*. In comparison to ODDA, the best performing heuristic requires on average 61.71% less PLF-C with a minimum of 1.99% and a maximum of 92.31%. The *extra_α_* of the best heuristic is surprisingly 0.81% lower than that of the ODDA. It might appear counter-intuitive, that some heuristics yield a lower *extra_α_* than the ODDA. However, our reshuffling strategies can strive to reduce *extra_α_* by placing entire small partitions with a large SRC onto the same CPU. Also, running ODDA on lexicographically sorted MSAs does not significantly change the results. It merely improves the PLF-C to 175.9% with respect to the lower bound *L*. Thus, a dedicated SR-aware data distribution heuristic *is* required since it can reduce PLF-C values and hence parallel runtimes by 60% on average.

The results of these performance analyses are summarized in Figure 12. The Figure displays the key heuristic performance metrics averaged over all MSAs. It also demonstrates that *extra_α_* is generally low and does not vary substantially among heuristics. Figure 8 presents a representative overview of the heuristic performances for dataset 2.

**Figure 8.**
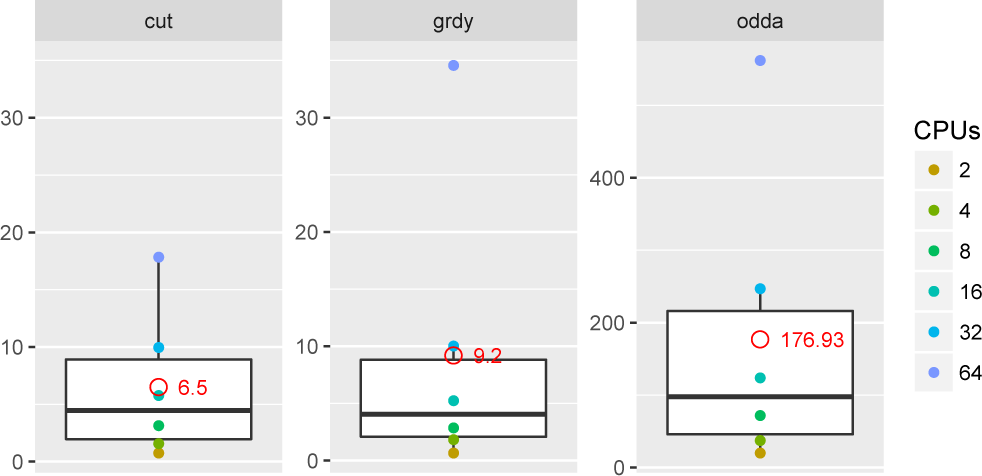
Comparison of the 2 best performing heuristic component combinations and the ODDA for dataset 2 for all 6 CPU scenarios. The y-axis shows the percentage that the most loaded CPU is larger than the lower bound L in terms of PLF-C. The red dot represents the average over all CPUs scenarios. In this case the heuristic using CUT requires on average 1 – (106.5/276.93) = 61.54% less PLF-C than the ODDA.

**Figure 9.**
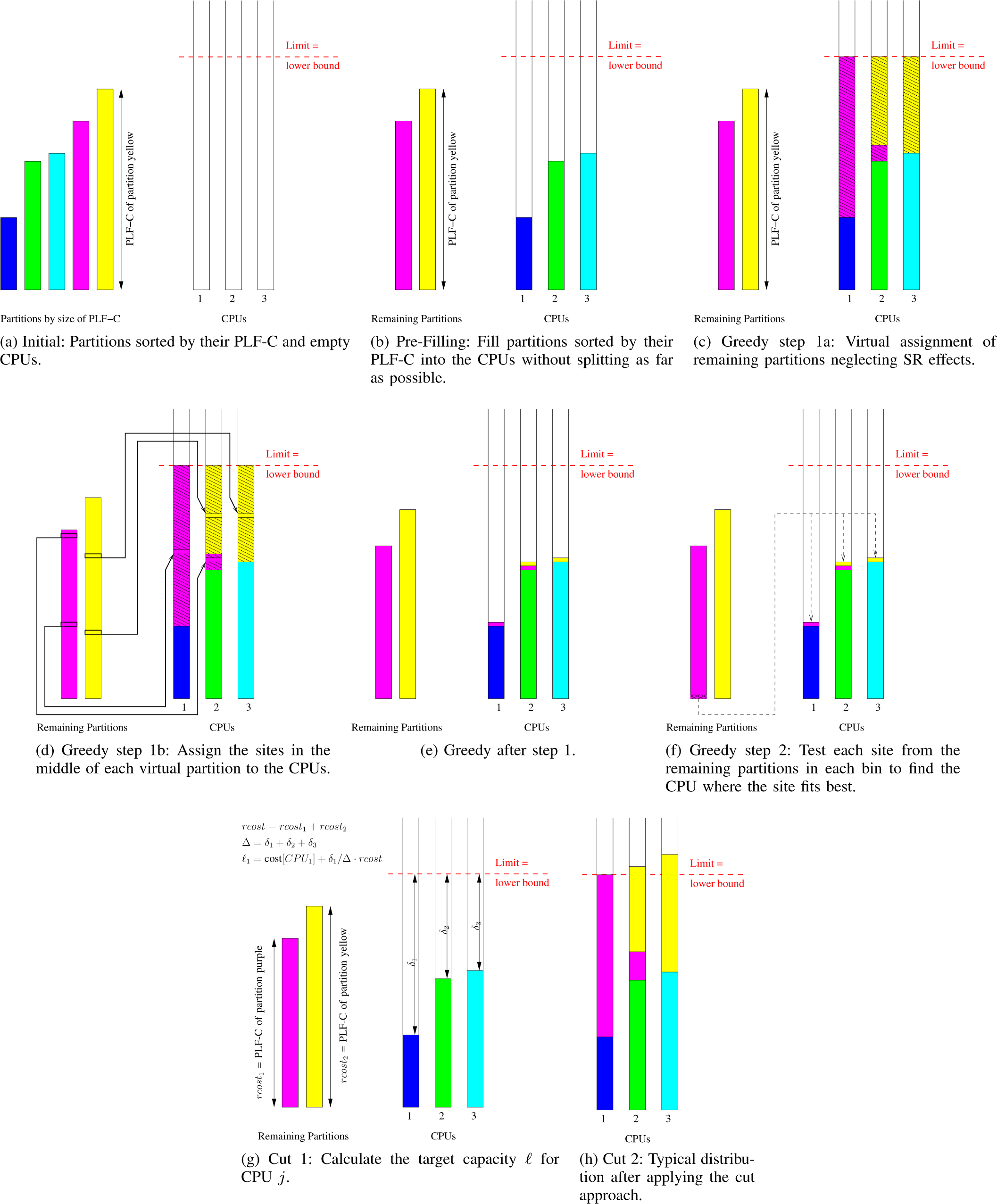
Illustrations.

In Figure 13 we provide an example data distribution for MSA 1 on 4 CPUs for our two best heuristics as well as for ODDA.

**Figure 13.**
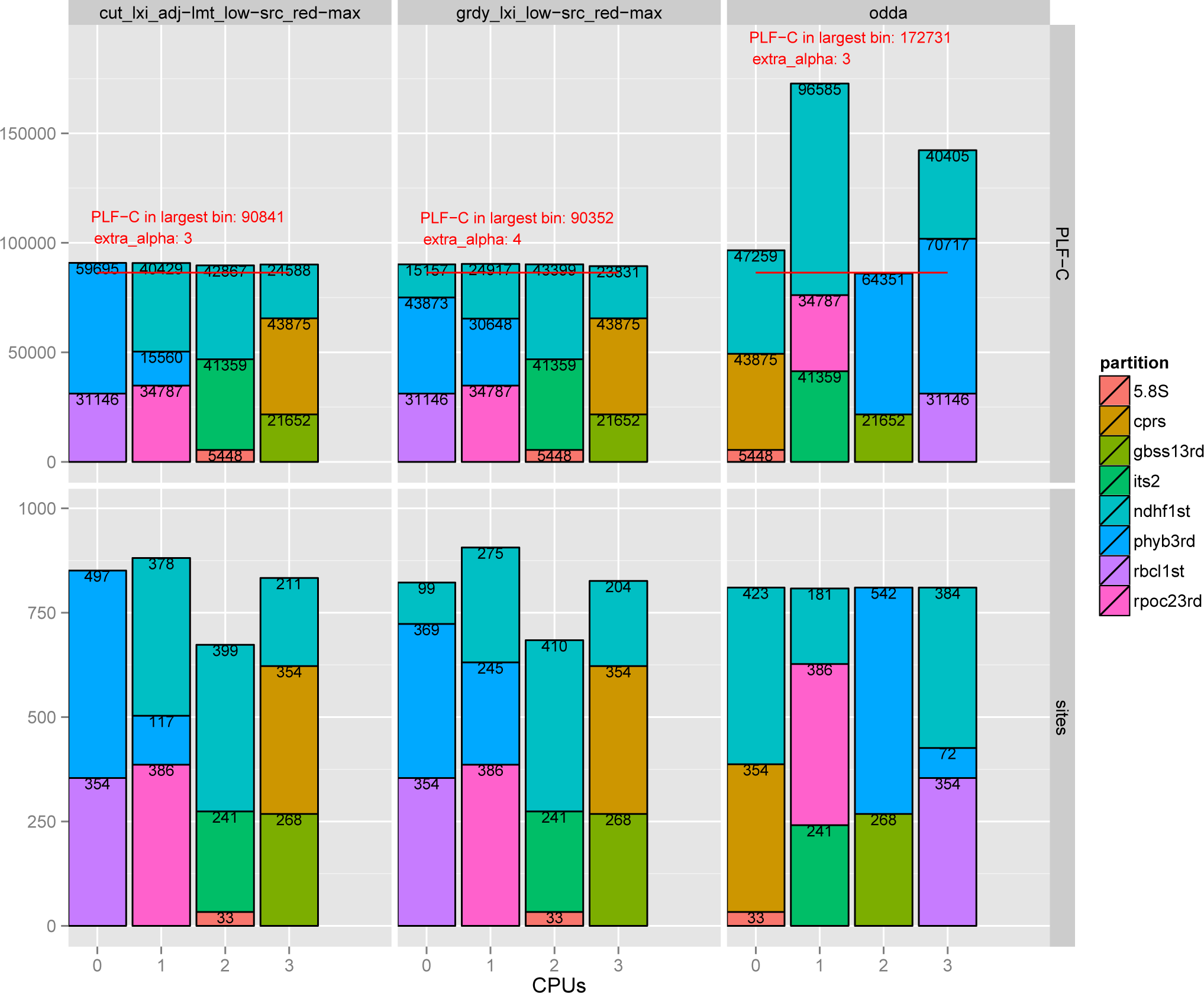
MSA 1 distributed across 4 CPUs. The upper part of the graph presents the data distribution in terms of PLF-C, the lower part in terms of sites. The red horizontal bar indicates the lower bound *L*. In this example the *greedy approach* in conjunction with a lexicographic MSA sort, the *low SRC* and *reduce max* reshuffling strategies performs best. The *extra_α_* value shows that one more redundant *P*(*t*) calculation is required than for the data distribution of the *cut approach.* The ODDA balances the partitions in terms of sites, but this deteriorates PLF-C values for SR-based PLF implementations.

## VII. CONCLUSION AND FUTURE WORK

In this paper we present the, to our knowledge, first data distribution problem statement for parallel PLF implementations that rely on the site repeats technique for reducing the number of PLF calculations on partitioned phylogenomic MSAs.

Whether the data distribution problem is NP-hard remains an open question. Here, we assume no polynomial algorithm exists, and we provide a lower bound *L* for the optimal data distribution scheme and explore a large number of heuristic data distribution algorithms using a component-based framework. All components and heuristics as well as the MSAs are freely available via `github`.

We initially characterize and quantify some aspects of SR-based calculations. Then, we present the two best performing heuristics that generate data distribution schemes with a PLF-C that is on average only 6% worse than the PLF-C of our conservative lower bound *L*. More importantly, we show that designing SR-aware data distribution algorithms *does matter*, since the standard SR-agnostic ODDA approach yields an average PLF-C for the most loaded CPU that is more than twice as high as the PLF-C attained by our heuristics. Thus, SR-aware data distribution heuristics can reduce runtimes for parallel phylogenetic analyses by 60%.

We are planning on integrating the heuristics into real-world phylogenetic inference tools. We are currently developing a revised version of the PLL [7] that will include a full SR implementation and a Message Passing Interface (MPI) parallelization. Hence, we will also integrate a variant of the heuristics presented here. Note that, these heuristics were assessed on fixed trees with fixed (midpoint) rootings. While this covers application scenarios where trees remain fixed and only statistical model parameters on the tree are being optimized (e.g., divergence time estimates or tests for positive selection) this is not practical for tree searches where the tree topologies and virtual root positions constantly change. To this end, we will need to test appropriate heuristics that yield “good” data distributions for a larger set of trees and virtual rootings. Evidently, there is a tradeoff between slightly increased PLF-C costs and frequent data re-distributions. While the asymptotic complexities of our heuristics might appear to be large, we expect them to be amortized by the substantially higher empirical cost of likelihood-based tree evaluations.

## ACKNOWLEDGMENT

Part of this work was financially supported by the Klaus Tschira Foundation.

## Footnotes

1 https://github.com/conscho/phylo_scheduling

2 https://github.com/jhbadger/Newick-ruby

